# PIRCHE application versions 3 and 4 lead to equivalent T cell epitope mismatch scores in solid organ and stem cell transplantation modules

**DOI:** 10.1101/2024.01.09.574785

**Authors:** Benedict M Matern, Matthias Niemann

## Abstract

Elevated PIRCHE scores between recipient and donor in organ and stem cell transplantation have been shown to correlate with increased risk of donor-specific HLA antibodies and graft-versus-host disease, respectively. With each revision of the PIRCHE application server, it is critical to completely evaluate the predicted scores, and compare with previous revisions. This manuscript compares the newly introduced PIRCHE version 4.2 with its predecessor version 3.3, which has been widely used in retrospective studies, using a virtual cohort of 10,000 transplant pairs. In the stem cell transplantation module, both versions yield identical results for 100% of the test population. In the solid organ transplantation module, 97% of the test population has identical PIRCHE scores in both versions. The deviating cases could be attributed to a refinement in the PIRCHE algorithm’s specification. For the 3% of cases with deviations, the determined magnitude of the difference is likely to be below the detection limit for clinical effects. We hereby confirm the equivalence in PIRCHE scores generated by the application server versions 3.3 and 4.2.

## Introduction

Histocompatibility is a major concern impacting treatment success in allo-transplant domains such as stem cell transplantation (SCT) and solid organ transplantation (SOT).^1,2^ The Prediction of Indirectly ReCognizable Human leukocyte antigen Epitopes (PIRCHE) has been shown to be correlated with several post-transplant outcomes such as graft-versus-host disease or donor-specific HLA antibodies. The PIRCHE algorithm is implemented as a web application and accessible through www.pirche.com. The application aggregates a number of pre- and post-processing steps to validate input data, impute HLA genotyping data, extrapolating HLA sequence data and HLA peptide binding predictions. The current major application version 3 has been used by several groups and was validated in various settings (Table 1).

**Table 1:** Original research articles mostly considered PIRCHE versions 2.X and 3.X.

| Title | First Author | Journal | Year | PIRCHE Version |
| --- | --- | --- | --- | --- |
| Donor-Recipient Matching Based on Predicted Indirectly Recognizable HLA Epitopes Independently Predicts the Incidence of De Novo Donor-Specific HLA Antibodies Following Renal Transplantation | Lachmann et al<sup>13</sup> | American Journal of Kidney Diseases | 2017 | 2.4.21 |
| The clinical significance of epitope mismatch load in kidney transplantation: A multicentre study | Daniels et al<sup>14</sup> | Transplant Immunology | 2018 | 2.5 |
| PIRCHE-II Is Related to Graft Failure after Kidney Transplantation | Geneugelijk et al<sup>15</sup> | Frontiers in Immunology | 2018 | 2.4 |
| Predicted Indirectly ReCognizable HLA Epitopes (PIRCHE) Are Associated with Poorer Outcome after Single Mismatch Unrelated Donor Stem Cell Transplantation | Ayuk et al<sup>16</sup> | Transfusion Medicine and Hemotherapy | 2019 | 1.0 |
| Exploratory Study of Predicted Indirectly ReCognizable HLA Epitopes in Mismatched Hematopoietic Cell Transplantations | Geneugelijk et al<sup>17</sup> | Frontiers in Immunology | 2019 | 2.0 |
| Exploring predicted indirectly recognizable HLA epitopes (PIRCHE-II) in liver transplant recipients on calcineurin inhibitor-free maintenance immunosuppression. A retrospective single center study. | Meszaros et al<sup>18</sup> | Transplant Immunology | 2020 | 3.0 |
| Donor/Recipient HLA Molecular Mismatch Scores Predict Primary Humoral and Cellular Alloimmunity in Kidney Transplantation | Meneghini et al<sup>19</sup> | Frontiers in Immunology | 2021 | 3.3 |
| Intravenous immunoglobulins do not prove beneficial to reduce alloimmunity among kidney transplant recipients with BKV-associated nephropathy | Naef et al<sup>20</sup> | Transplant International | 2021 | 3.1 |
| Computational Eurotransplant kidney allocation simulations demonstrate the feasibility and benefit of T-cell epitope matching | Niemann et al<sup>21</sup> | PLoS Computational Biology | 2021 | 3.1 |
| Peptides Derived From Mismatched Paternal Human Leukocyte Antigen Predicted to Be Presented by HLA-DRB1, -DRB3/4/5, -DQ, and -DP Induce Child-Specific Antibodies in Pregnant Women | Niemann et al<sup>22</sup> | Frontiers in Immunology | 2021 | 3.2 |

*“PIRCHE application versions 3 and 4 lead to equivalent T cell epitope mismatch scores in solid organ and stem cell transplantation modules”;* Ben Matern, Matthias Niemann; © 2024 PIRCHE AG

|  |  |  |  |  |
| --- | --- | --- | --- | --- |
| T-Cell Epitopes Shared Between Immunizing HLA and Donor HLA Associate With Graft Failure After Kidney Transplantation | Peereboom et al<sup>23</sup> | Frontiers in Immunology | 2021 | 3.3.30 |
| Risk factors, histopathological features, and graft outcome of transplant glomerulopathy in the absence of donor-specific HLA antibodies | Senev et al<sup>24</sup> | Kidney International | 2021 | 3.1 |
| Clinical Significance of Shared T Cell Epitope Analysis in Early De Novo Donor-Specific Anti-HLA Antibody Production After Kidney Transplantation and Comparison With Shared B cell Epitope Analysis | Tomosugi et al<sup>25</sup> | Frontiers in Immunology | 2021 | 3.0 |
| Can PIRCHE-II Matching Outmatch Traditional HLA Matching? | Unterrainer et al<sup>26</sup> | Frontiers in Immunology | 2021 | 3.0 |
| Non-invasive alloimmune risk stratification of long-term liver transplant recipients | Vionnet et al<sup>27</sup> | Journal of Hepatology | 2021 | 3.3.26 |
| The number of donor HLA-derived T cell epitopes available for indirect antigen presentation determines the risk for vascular rejection after kidney transplantation | Betjes et al<sup>28</sup> | Frontiers in Immunology | 2022 | 3.3.43 |
| Dissecting the impact of molecular T-cell HLA mismatches in kidney transplant failure: A retrospective cohort study | Lemieux et al<sup>12</sup> | Frontiers in Immunology | 2022 | 3.2 |
| High PIRCHE Scores May Allow Risk Stratification of Borderline Rejection in Kidney Transplant Recipients | Lezoeva et al<sup>29</sup> | Frontiers in Immunology | 2022 | 3.3 |
| Formation of donor-specific antibodies depends on the epitope load of mismatched HLAs in lung transplant recipients: A retrospective single-center study | Lobashevsky et al<sup>30</sup> | Clinical Transplant | 2022 | 3.1 |
| Immunologic Risk Stratification of Pediatric Heart Transplant Patients by Combining HlaMatchmaker and PIRCHE-II | Mangiola et al<sup>31</sup> | The Journal of Heart and Lung Transplantation | 2022 | 3.3.55 |
| Snowflake epitope matching correlates with child-specific antibodies during pregnancy and donor-specific antibodies after kidney transplantation | Niemann et al<sup>32</sup> | Frontiers in Immunology | 2022 | 3.3 |
| Analysis of T and B Cell Epitopes to Predict the Risk of de novo Donor-Specific Antibody (DSA) Production After Kidney Transplantation: A Two-Center Retrospective Cohort Study | Sakamoto et al<sup>33</sup> | Frontiers in Immunology | 2022 | 3 |

*“PIRCHE application versions 3 and 4 lead to equivalent T cell epitope mismatch scores in solid organ and stem cell transplantation modules”;* Ben Matern, Matthias Niemann; © 2024 PIRCHE AG

|  |  |  |  |  |
| --- | --- | --- | --- | --- |
| PIRCHE-II scores prove useful as a predictive biomarker among kidney transplant recipients with rejection: An analysis of indication and follow-up biopsies | Spitznagel et al<sup>34</sup> | Frontiers in Immunology | 2022 | 3.3 |
| Immunologic risk stratification of pediatric heart transplant patients by combining HLA-EMMA and PIRCHE-II | Ellison et al<sup>35</sup> | Frontiers in Immunology | 2023 | 3.3.57 |
| Predictive value of molecular matching tools for the development of donor specific HLA-antibodies in patients undergoing lung transplantation | Kleid et al<sup>36</sup> | HLA Immune Response Genetics | 2023 | 3.3 |

To extend the application to new transplant domains, incorporate proteins beyond classical HLA, and support implementation of new features, the internal data structures and software architecture of the PIRCHE application server were redesigned yielding the new major application version 4. Using test-first development, regression tests have been implemented prior to software design changes.

The PIRCHE algorithm has been described in 2013 by a Dutch research group of Eric Spierings in the context of kidney transplantation.^3^ Shortly after, the definition has been adapted to stem cell transplant domains.^4^ In brief, the PIRCHE-II score in SOT is defined as the number of unique donor-HLA derived allo-peptides predicted to be presented by recipient Class II HLA, that are not present in the presentable self-peptidome. Conversely in SCT, PIRCHE are defined as recipient-HLA derived allo-peptides presented by matched HLA Class I or Class II to reflect graft-versus-host responses. Several constraints have been defined to allow for consistent calculation of numbers. An updated and more refined formal definition of the PIRCHE score in SOT has been provided by Niemann et al.^5^ It refines the definition of the presentable self-peptidome by restricting it to peptides derived from extracellular parts of the HLA proteins by excluding the leader peptide. This updated specification has been followed in the implementation of PIRCHE version 4.

Within this study, we’re calculating PIRCHE scores of 10,000 virtual transplant pairs generated through the application server versions 3.3 and 4.2 considering an identical IMGT database (3.47)^6^, HLA cleavage predictor^7^, HLA peptide binding predictors^8,9^ and genotype imputation pipeline^10^. The results of both versions are compared to identify the frequency of cases with deviating scores and quantify the magnitude of these deviations.

## Materials and Methods

Patient and donor populations were simulated based on an equal distribution of the populations in the 2007 National Marrow Donor Program frequency tables.^11^ These frequencies distinguish four ethnicities (i.e. African Americans, Asians and Pacific Islanders, European Americans, and Hispanics), and 1,250 were selected from each to create a dataset of 5,000 individuals for both the donor and patient populations. This provides reasonably distributed simulated populations over varied ethnicities.

The 5,000 patients and 5,000 donors were shuffled and randomly assigned to each other to simulate paired SOT. By shuffling and randomly pairing the simulated populations, the dataset represents transplantations both within an ethnicity group, as well as across ethnicities.

For SCT, the analysis was performed on a simulated 9/10 HLA-matched dataset. To this end, for each of the 5,000 patient genotypes, one of the 10 alleles from HLA-A, -B, -C, -DRB1, or -DQB1 was randomly selected. This allele is replaced with a random allele from the same locus, which was not present in the patients’ own genotype.

The pairings were written to a CSV file in the PIRCHE bulk analysis format containing 5,000 transplantations for both the SOT and SCT test dataset, respectively. These were submitted to the corresponding PIRCHE bulk CSV analysis endpoints using both the current version of PIRCHE (3.3) and the new version (4.2) in the SOT and SCT modules. The PIRCHE server was configured using the default thresholds of 0.5 for cleavage, 500 nM for Class I binding affinity and 1000 nM for Class II binding affinity. Results were temporarily stored remotely on the PIRCHE server. The bulk CSV module returns PIRCHE scores and corresponding allo-peptides for each pairing, which were then exported, parsed and compared.

PIRCHE scores between the versions were compared (SCT: PIRCHE-I/-II, SOT: PIRCHE-II), as well as the natural log-transformed one-incremented scores as suggested by Geneugelijk et al.^10^ In the cases where scores between versions differed, the details of the PIRCHE peptides (presenting protein and 9-mer core peptide) were identified. Core peptides were further mapped to their source 15-mer peptide and the corresponding source HLA protein. This resulted in a categorization of exactly which peptides were discrepant between versions.

## Results

The cohorts considered a total of 395 distinct HLA proteins (HLA-A: 84, HLA-B: 154, HLA-C: 62, HLA-DRB1: 73, HLA-DQB1: 22). The distribution of PIRCHE scores is similar between the two versions. Summary statistics for the score ranges for the simulated test panel were identical between versions 3.3 and 4.2. (Table 2)

**Table 2:** PIRCHE score summary statistics for simulated test panel.

|  | Min | Min (Ln) | Median | Med (Ln) | Max | Max(Ln) |
| --- | --- | --- | --- | --- | --- | --- |
| **SOT**<br>**PIRCHE-II** | 5 | 1.61 | 89 | 4.49 | 316 | 5.76 |
| **SCT**<br>**PIRCHE-I** | 0 | 0 | 4 | 1.39 | 35 | 3.56 |
| **SCT**<br>**PIRCHE-II** | 0 | 0 | 10 | 2.30 | 78 | 4.36 |

On case-by-case comparisons, scores showed good agreement. In nearly all cases, PIRCHE scores were identical. In SCT, we found both versions yielding congruent results. All cases had the same PIRCHE scores in the two algorithm versions. (Figures 1 and 2)

**Figure 1:**
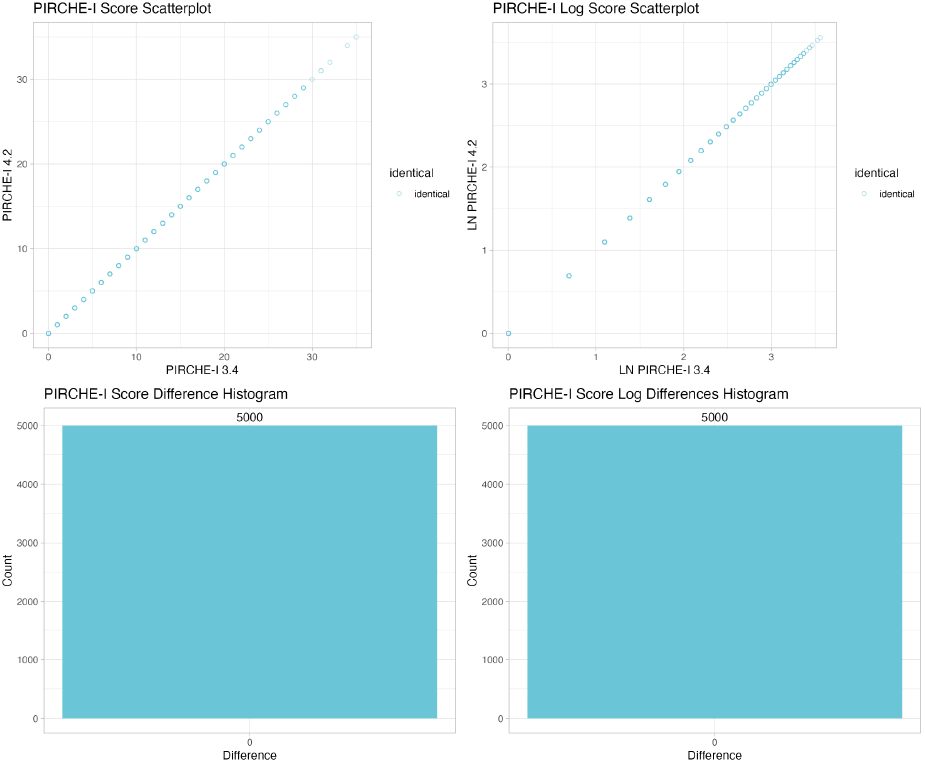
Comparison of PIRCHE-I scores in the SCT context. No differences between the versions have been observed.

**Figure 2:**
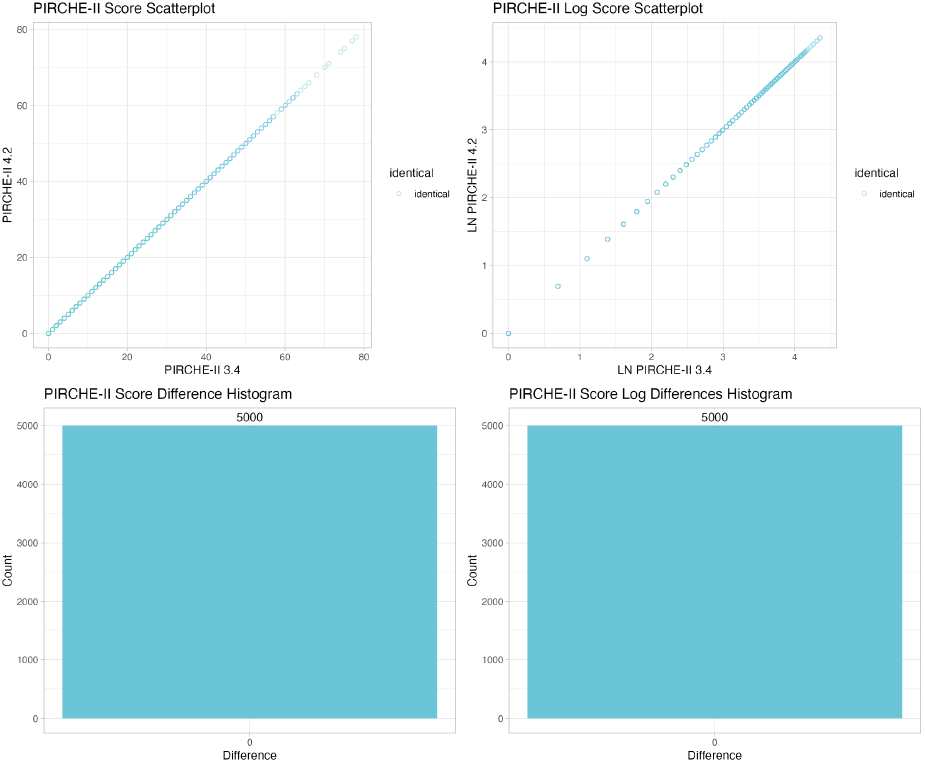
Comparison of PIRCHE-II scores in the SCT context. No differences between the versions have been observed.

In SOT, however, we identified a few differences. (Figure 3) In 4845 (96.9%) the scores were identical. In 150 (3%) we identified off-by-one scores. In 5 (0.1%) cases, the score difference was 2. In all of these discrepancies, the numbers of allo-peptides calculated by version 4.2 were higher than in the previous version, indicating one or two extra PIRCHE peptides were found in the newer version.

**Figure 3:**
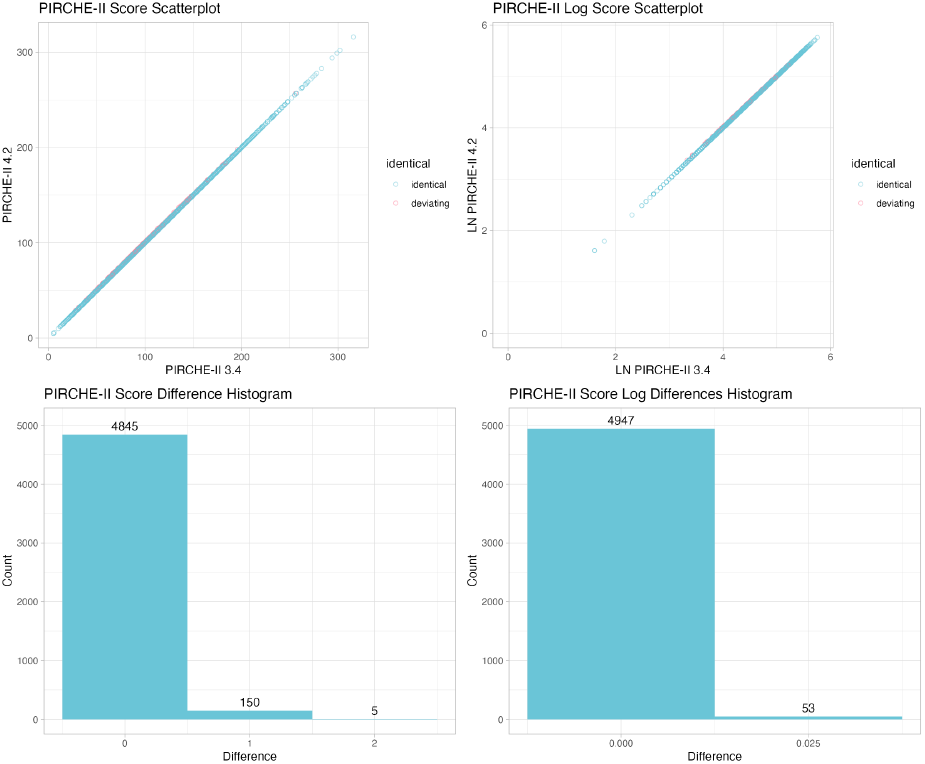
Comparison of PIRCHE-II SOT scores. In 3.1% of cases, 1-2 extra PIRCHE peptides were discovered in the 4.2 version. Differences in natural logarithm PIRCHE scores are very low.

Comparisons of differences in the log-transformed PIRCHE-II scores show very miniscule differences. In our analysis we observed a maximum difference of 0.035. The score discrepancies tend to occur in the cases of higher scores, where clinical relevance of differences in one peptide are small compared to the score magnitude.

Investigations of these score differences show distinct patterns in the peptides responsible for the discrepancies (Table 3). In this data set, the discrepancies were caused by three distinct core peptides.

**Table 3:** Additional PIRCHE core peptides identified in the SOT dataset comparison.

| Peptide Core | 15mer peptides | Source | Presenter | # Cases |
| --- | --- | --- | --- | --- |
| MRYFSTSVS | SHSMRYFSTSVSWPG | C*04:01 | DRB1*07:01 | 79 |
| MRYFSTSVS | SHSMRYFSTSVSWPG | C*04:01 | DRB1*09:01 | 34 |
| MRYFYTAVS | CSHSMRYFYTAVSRP | C*05:01 | DRB1*14:03 | 1 |
| MRYFYTAVS | CSHSMRYFYTAVSRP | C*12:02 | DRB1*14:03 | 1 |
| MRYFYTAVS | CSHSMRYFYTAVSRP | C*15:02 | DRB1*14:03 | 1 |
| MRYFYTAVS | CSHSMRYFYTAVSRP | C*16:01 | DRB1*14:03 | 1 |
| SMRYFSTSV | CSHSMRYFSTSVSRP | C*14:02 | DRB1*13:12 | 1 |
| SMRYFSTSV | CSHSMRYFSTSVSRP | C*14:02 | DRB1*14:04 | 2 |
| SMRYFSTSV | CSHSMRYFSTSVSRP | C*14:02 | DRB1*15:03 | 3 |
| SMRYFSTSV | GSHSMRYFSTSVSRP | A*23:01 | DRB1*15:02 | 2 |
| SMRYFSTSV | GSHSMRYFSTSVSRP | A*23:01 | DRB1*15:03 | 6 |
| SMRYFSTSV | GSHSMRYFSTSVSRP | A*24:02 | DRB1*15:02 | 7 |
| SMRYFSTSV | GSHSMRYFSTSVSRP | A*24:02 | DRB1*15:03 | 11 |
| SMRYFSTSV | GSHSMRYFSTSVSRP | A*30:01 | DRB1*15:02 | 3 |
| SMRYFSTSV | GSHSMRYFSTSVSRP | A*30:01 | DRB1*15:03 | 4 |
| SMRYFSTSV | GSHSMRYFSTSVSRP | A*30:02 | DRB1*15:03 | 6 |
| SMRYFSTSV | **SMRYFSTSVSWPGRG** | C*04:01 | DRB1*13:12 | 5 |

## Discussion

This is the first study to quantitatively and qualitatively compare differences between the new PIRCHE application major version 4 and the current version 3. Previously Lemieux et al reported PIRCHE score deviations when considering the identical application server version yet considering a different version of the peptide binding prediction pipeline.^12^ In that study, an improved algorithm to complete non-sequenced regions of reported HLA proteins has been introduced, causing discrepancies between results.

Our data revealed identical results between both versions considering the PIRCHE SCT matching modules. This confirms the new application server version follows the specification which - for SCT - remained identical for both versions 3 and 4. For the SOT matching modules we experienced increased scores occasionally, which follows a slight change in the specification for the SOT algorithm.

Through application debugging of version 3 considering a representative example case based on Table 3, the deviation in the considered self-peptidome has been confirmed as the underlying reason for discrepant results. The specification of the version 4 SOT algorithm considers only HLA-presented core peptides located on the extracellular domains of the donor HLA as self-peptidome. Version 3 conversely considers the self-peptidome also based on intracellular protein fragments. This increased the self-peptidome in version 3 resulting in a slightly lower average PIRCHE-II score that manifests in rare cases, where differences in a core peptide’s flanking region located in exon 1 differ the preference for a binding core. Consequently in SOT, PIRCHE version 4 may yield a slightly increased score in a few cases. Notably this effect is not apparent in the PIRCHE SCT algorithms, given the specification in both versions 3 and 4 consider the full protein including exon 1 both for the self-peptidome and the mismatch-derived peptidome.

In a previous study on multiple HLA genotype imputation by Geneugelijk et al,^10^ a delta log score of +/-1 was considered as a high deviation, given the clinically relevant PIRCHE score ranges as suggested by Lachmann et al, that were based on log-transformed PIRCHE scores.^13^ The greatest observed delta log score of +0.035 in the SOT comparison is way below these score deviations, confirming the negligible clinical impact of the deviation identified in both application versions. This suggests that also in the most deviant cases, the difference is not likely to lead to differences in interpretation of graft failure likelihood or other clinical relevance. For ∼97% of the cases, congruence has been shown.

Underrepresentation of rare alleles potentially limits generalizability of our study results. The applied haplotype frequency-based approach of generating a virtual transplant population prefers frequent haplotypes and alleles. Even though we are using different ethnicities, we only consider 395 of the currently over 18,000 reported proteins based on HLA-A, -B, -C, -DRB1 and -DQB1 alleles. Alternative approaches could use more refined haplotype frequencies or virtual populations irrespective of allele frequencies. Furthermore, restricting the validation to 5 HLA loci may miss discrepancies introduced through other loci. Given our validation however could confirm the underlying reason for discrepancies is a slight refinement of the specification, the overall impact on score deviations remains very limited, as mostly a potential core peptide’s flanking region exceeding into exon 1 changes the self-peptidome, whereas further leader peptide-derived peptides are of low relevance due to lacking peptide overlap with other exons.

The study design considered randomly paired organ transplants, and randomly simulated 9/10 transplants. This approach overestimates the average PIRCHE scores seen in transplantation, given HLA-based allocation is widely applied. Consequently it does not fully represent the PIRCHE scores that may be observed in a real panel of transplantations. As such the PIRCHE score distributions as reported herein are not comparable to PIRCHE scores reported by published studies.

As presented in Table 1, early studies on PIRCHE have also considered the major application versions 1 and 2. Notably, these versions did not introduce major changes to the implemented matching algorithm, but added features outside the matching program logic, whilst regression tests were in place.

In summary, we confirm the PIRCHE application versions 3 and 4 generating equivalent scores in the SCT and SOT matching modules.

## Acronyms

PIRCHE: Predicted Indirectly Recognizable HLA Epitopes
SCT: Stem Cell Transplantation
SOT: Solid Organ Transplantation

